# Tensor Kernel Learning for Classification of Alzheimer’s Conditions using Multimodal Data

**DOI:** 10.1101/2024.07.17.603875

**Authors:** Vu Duy Thanh, Thanh Trung Le, Pham Minh Tuan, Nguyen Linh Trung, Karim Abed-Meraim, Mouloud Adel, Nguyen Viet Dung, Nguyen Thanh Trung, Dinh Doan Long, Oliver Y. Chén

**Author notes:** Corresponding author: Nguyen Linh Trung.

## Abstract

Early, timely, and accurate assessment of Alzheimer’s disease (AD), particularly at its earlier stage – mild cognitive impairment (MCI) –, is central to detecting, managing, and potentially treating the disease. The biological underpinnings of AD, however, is multifaceted, from genetic variations, abnormal protein accumulation, to irregular brain functions and structure. A joint analysis of these data, therefore, may offer potentially new insights about AD-related biomarkers and AD prediction. But such explorations must confront the complexity of these data: heterogeneity, multimodality and high-dimensionality. Here, to address these challenges, we propose a new machine-learning method, namely *the tensor kernel learning (TKL)*, leveraging tensor methods and kernel learning, to enhance AD assessment by enhancing multi-modal data integration. More specifically, TKL first uses CP/PARAFAC decomposition and graph diffusion to fuse multiple kernels learned from four complementary data modalities (MRI, PET, CSF, and SNP data). We then used a supervised kernel for a kernel SVM classifier to identify potential patients. To evaluate the effectiveness of TKL, we apply it to data from the Alzheimer’s Disease Neuroimaging Initiative (ADNI; the sample size *n* = 331 subjects), including cognitively normal individuals (CN), MCI subjects, and AD patients. TKL improves AD classification performance in both linear and nonlinear combinations, achieving accuracies of 91.31% for CN vs. AD, 81.45% for CN vs. MCI, and 78.27% for AD vs. MCI, compared to 85.48%, 70.89%, and 73.51% using the best single modality. Additionally, TKL reveals clearer, more structured patterns in the data, enhancing interpretation and understanding of the relationships among different modalities. Detailed implementation details of our method can be found at: https://github.com/thanhvd18/Tensor-Kernel-Learning-matlab.

## I. Introduction

Alzheimer’s disease (AD) is a progressive and prevalent neurodegenerative disorder primarily affecting older adults. It is characterized by the accumulation of beta-amyloid and tau, proteins that lead to cell death and the obstruction of transportation inside neurons [1]. AD poses significant health, medical, and societal challenges [2]: it affects approximately 17% of people aged 65 and above [3], [4] and the number of people suffering from AD worldwide is expected to reach 152 million by 2050. Unfortunately, there is, as of yet, no cure for AD [5]. Thus, the early and timely assessment of AD may be crucial to help better manage the disease.

Chief to AD assessment is the discovery of AD-related biomarkers. Biomarkers play a crucial role in detection, assessing, and monitoring of disease progression, particularly in its early stages. Common data for AD biomarker development include magnetic resonance imaging (MRI), positron emission tomography (PET), cerebrospinal fluid (CSF), and single nucleotide polymorphisms (SNPs) data [6]. Among them, MRI is perhaps one of the most widely used in studying dementia diseases in general and AD in particular [7]. Examples of MRI features are cerebral atrophy (or ventricular expansion), voxel-wise tissue analysis, cortical thickness, and volume, among others. Fluorodeoxyglucose positron emission tomography (FDG-PET) has also been used in AD studies [8], [9]. For example, FDG-PET imaging enhances diagnostic accuracy by unveiling glucose metabolism reduction in specific brain regions among subjects with AD [10]. Apart from neuroimaging modalities, cerebrospinal fluid (CSF) biomarkers, such as A*β*_42_, total tau (T-tau), and phosphorylated tau (P-tau), provide valuable information for AD diagnosis [11]. For example, AD subjects generally display low values of A*β*_42_ and high concentrations of T-tau and P-tau. Finally, genetic factors, notably the apolipoprotein E (APOE) gene, constitute significant risk factor to AD [1], [3].

Together, these lines of evidence suggest that different data modalities may reveal distinct aspects of pathological changes associated with AD and offer converging or complementary diagnostic insights. To constructively combine information from multimodal data to better inform AD assessment and prediction, one, however, needs effective data integration and fusion methods coupled with potent predictive models [12].

Much effort has been made to develop multimodal data fusion methods to, in part, address the limitations of single-modality-based methods in disease prediction [12], [13]. Broadly, these methods can be categorized into four groups: (i) correlation-based, (ii) cluster-based, (iii) multitask learning-based, and (iv) multiple kernel learning-based methods. In brief, correlation-based methods, such as [14], [15], are useful for unveiling shared patterns among modalities. Clustering-based methods find shared clusters among modalities [16], [17]. . Fusion methods based on multitask learning utilize labeled modalities, translating fusion challenges into relatively easier multitask learning problems. The optimal, shared multimodal features are selected by maximizing the classification accuracy [18], [19]. The last approach leverages multiple kernel (or multi-kernel) learning techniques and has gained attention for its capacity to integrate diverse sources of information [20]. By combining kernels derived from different data sources, it accommodates and integrates a diverse range of heterogeneous information. Thus, multiple kernel learning is particularly advantageous when dealing with heterogeneous data types and helps build more robust and accurate models. Consequently, studying multi-modal data with multi-kernel learning not only improves predictive accuracy, but also, by comparing the modality-specific kernels, helps better understand the complex biological systems [21]. Thanks to these attractive properties, we adopt the concept of multiple kernel learning, and combine it with tensor decomposition (see below), for treating the AD classification/detection problem using multi-modal data.

Recent years have seen successful uses of tensor decomposition (TD) in computer vision [22], [23], genetics [24], [25], and source separation [26], [27] (see [28], [29] for comprehensive surveys). In tensor fusion, a common approach involves concatenating multiple data tensors into a single one and employing TD to obtain shared features [30], [31]. While TD enhances model performance, simplifies parameter structures, and improves interpretation, challenges arise when dealing with multi-modal heterogeneous, large-scale, and high-dimensional data. The concatenated tensor often grows exponentially, posing significant difficulties and huge variability for current TD methods. Even if decomposition is feasible, the resulting components (e.g., core tensors and loading factors) may not be optimal, affecting the quality of the shared features.

To address this, we propose *Tensor Kernel Learning (TKL)*, which fuses heterogeneous data into a common supervised kernel space. TKL thus not only fuses multimodal data in the implicit feature space, avoiding the need for large or computationally expensive original features, but also takes advantage of the structural information among modalities. We employed directly supervised kernels for classification instead of adding manifold learning to obtain low-dimensional representations as done in [32], [33]. Experimentally, our method integrated well with both linear and nonlinear combinations for AD classification tasks using multi-modal data.

## II. Materials and Proposed Method

### A. Materials

#### 1) Dataset

This paper considers 331 subjects from the Alzheimer’s Disease Neuroimaging Initiative (ADNI) data (www.loni.ucla.edu/ADNI): 121 subjects who are cognitively normal, 100 with Mild Cognitive Impairment (MCI), and 110 with AD. See Table I for detailed demographic information.

**TABLE I.** Demographic by group for the studied sample. nO. = Number of subjects. Age = Mean age (± standard deviation).

|  | CN |  | MCI |  | AD |  |
| --- | --- | --- | --- | --- | --- | --- |
|  | No. | Age | No. | Age | No. | Age |
| Male | 66 | 74.62 ± 5.60 | 67 | 76.39 ± 6.70 | 68 | 75.47 ± 7.90 |
| Female | 55 | 74.71 ± 5.33 | 33 | 72.25 ± 7.54 | 42 | 72.61 ± 7.53 |
| Total | 121 | 74.66 ± 5.46 | 100 | 75.03 ± 7.23 | 110 | 74.37 ± 7.85 |

The neuroimaging data consist of T1-weighted MRI and FDG-PET images. See Fig. 1 for example images from AD and CN subjects. The MRI images contain 176 × 240 × 256 voxels (sized 1.2 × 1 × 1 mm^3^); each image is segmented into grey matter (GM), white matter, and CSF, with GM selected as the feature of interest. PET images comprise 160 × 160 × 96 voxels (sized 1.5 × 1.5 × 1.5 mm^3^). MRI highlights brain atrophy in specific regions; PET highlights reduced glucose metabolism in areas potentially affected by AD. We utilized three measurements for CSF features: A*β*42, PTAU, and TAU (see Fig. 2). For genetic information, we obtained single nucleotide polymorphisms (SNPs) data. We obtained the count of minor variants from ADNI (see Fig. 2) and removed SNPs with missing values, resulting in 6244 SNPs.

**Fig. 1.**
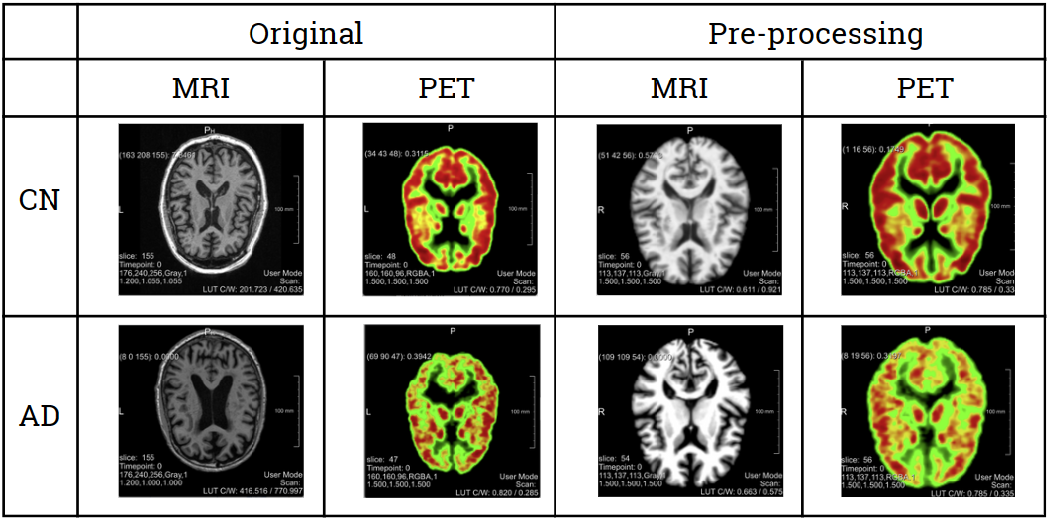
Brain imaging before and after preprocessing. The top row presents a slice of T1-weighted MRI alongside a corresponding slice of FDG-PET from a cognitively normal subject, while the bottom row depicts images from a subject diagnosed with AD.

**Fig. 2.**
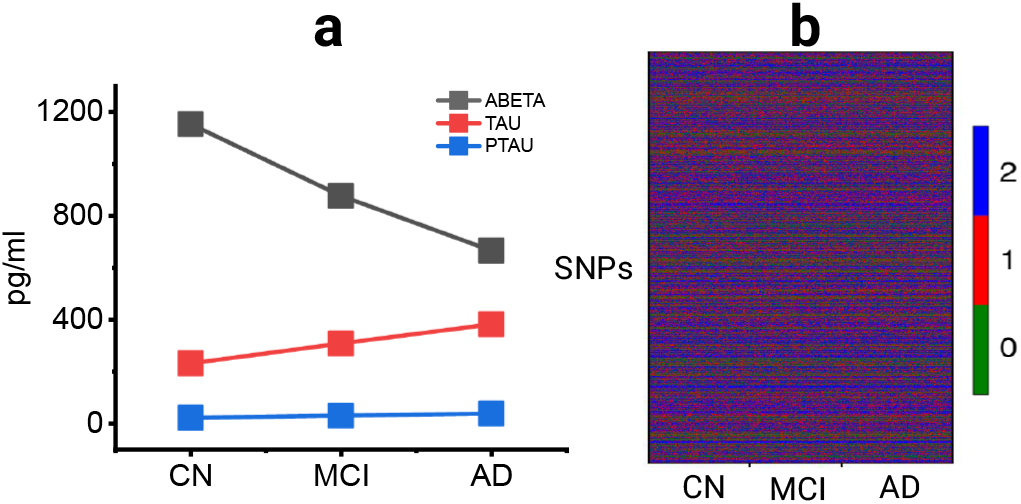
(a) Cerebrospinal fluid (CSF) data in this study. Patients with AD have abnormally low levels of A*β*42 and high levels of T-tau and P-tau in comparison with normal subjects. (b) An overview of SNPs data in this study. The rows represent 6244 SNPs and the columns are 331 subjects arranged by three groups. Each SNP indicates gene variation, represented as 0 for no major variant, 1 for one major variant, and 2 for two major variants.

#### 2) Pre-processing

The T1 MRI data underwent imaging by ADNI (https://adni.loni.usc.edu/methods/mri-tool/mri-pre-processing/). The pre-processed MRI images were then segmented to extract GM tissues using the CAT12 toolbox (https://neuro-jena.github.io/cat/). PET images were co-registered with MRI images using the ANTs toolbox (https://github.com/ANTsX/ANTs). Following the preprocessing procedures, all images were brought into the same MNI space, ensuring alignment both within the same subject across different modalities and across different subjects. We employed the template provided by the CAT12 toolbox, which has dimensions of 113 × 137 × 113 with isotropic voxels of 1.5 mm. Finally, both MRI and PET images were projected into the Schaefer 2018 atlas with 200 parcellations [34].

### B. The proposed Method

We present schematic pipeline of the proposed method in Fig. 3. We used four modalities: grey matter, FDG-PET, CSF and SNP. Each modality was independently trained by a random forest classifier and we obtained a similarity matrix for each modality. The four similarity matrices were then stacked into a three-way tensor. During tensor kernel learning, we approximated the tensor with a rank equal to the number of classes. Next, the similarity matrices exchanged information through a diffusion process and were combined into a single joint similarity matrix. This matrix was afterward fed into a kernel support vector machine (SVM), where the training and testing indices remained the same as in the random forest classifier step. Throughout, we used the term “similarity matrix” (output of the random forest) interchangeably with “kernel matrix” in kernel SVM.

**Fig. 3.**
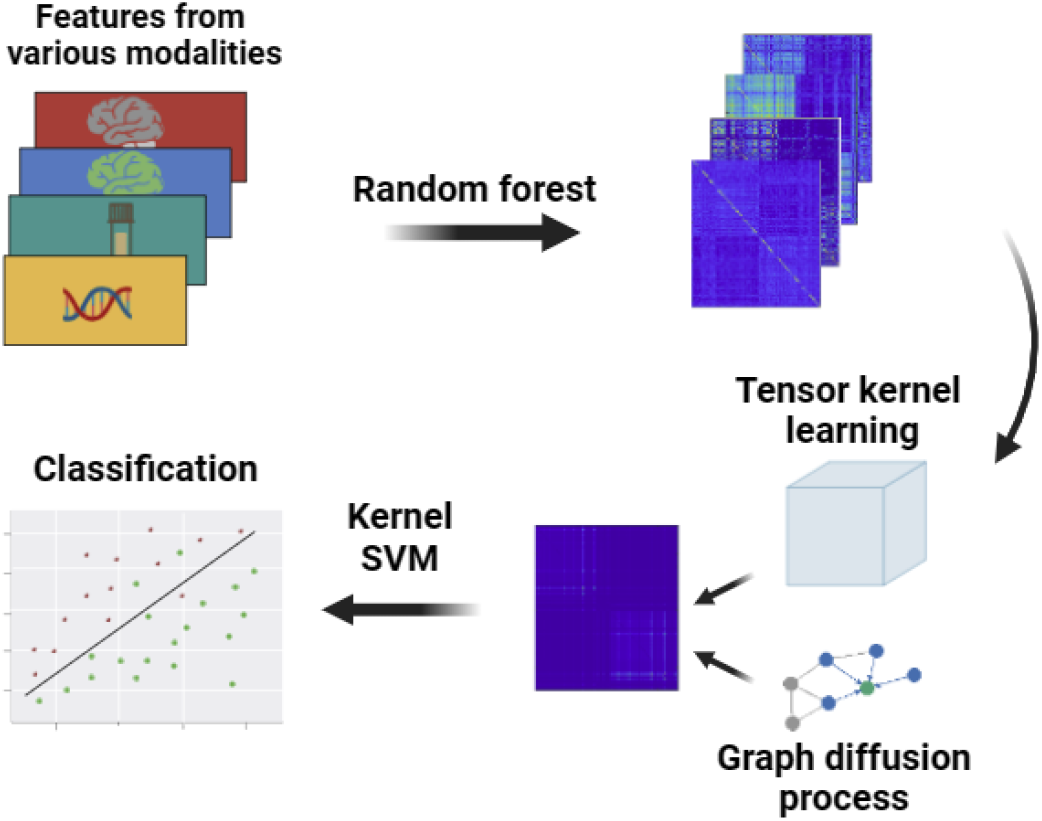
Overview of the framework and proposed method.

#### 1) Random forest classifier

We used a random forest (RF) classifier to generate the similarity matrix, following [32] (via *scikit-learn* library in Python). We chose 500 trees for each modality. The outcome is a similarity matrix among *N* subjects for each modality. The similarity between two subjects is determined by the proportion of trees in which the subjects end up in the same leaf. Additionally, we examined feature importance in the classification task within the RF. For example, we projected the identified features from MRI and PET data, respectively, onto the brain space in Fig. 4, where the colour code indicates feature importance. Our results suggest that PET features in certain areas are more important than their MRI counterparts. This is particularly pronounced in the temporal and parietal lobes, critical in AD [9]. Additionally, there seems little overlapping regions between MRI modality and PET modality. This hints that RF identified likely complementing features; with added non-overlapping information, this may potentially improve classification.

**Fig. 4.**
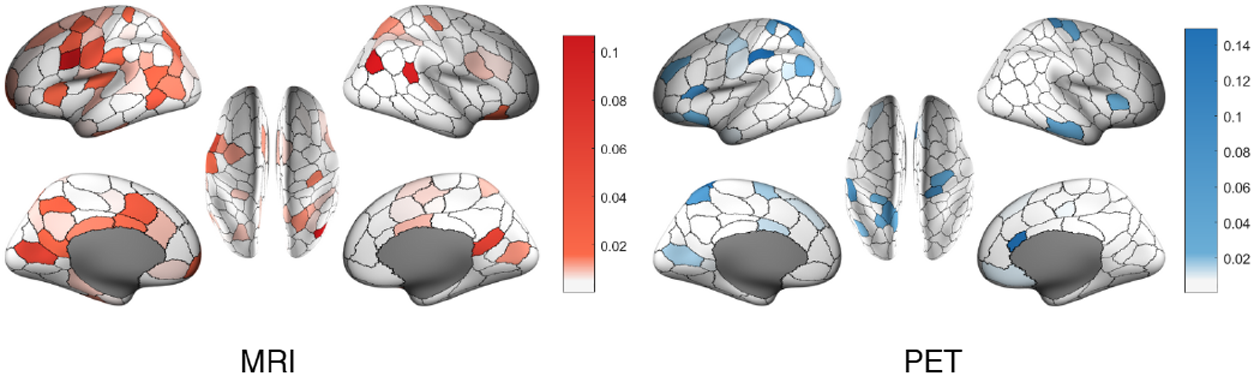
Identified features from MRI and PET data and their feature importance for distinguishing AD and CN.

#### 2) Tensor kernel learning (TKL)

After obtaining *M* similarity matrices (sized *N* × *N*) from *M* modalities, we concatenated them into a three-way tensor 𝒳 ∈ ℝ^*N*×*N*×*M*^. The objective was to estimate a low-rank tensor 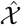 through CP/PARAFAC decomposition. Specifically, we found a low-rank tensor 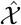 via the following optimisation problem:

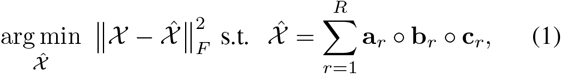

where “ ° “ denotes the outer product, **A** = [**a**_1_, **a**_2_, …, **a**_*R*_] ∈ ℝ^*N*×*R*^, **B** = [**b**_1_, **b**_2_, …, **b**_*R*_] ∈ ℝ^*N*×*R*^, and **C** = [**c**_1_, **c**_2_, …, **c**_*R*_] ∈ ℝ^*M*×*R*^ are loading factors. We set *R* as the number of classes in the multimodal AD classification task. To solve Equation (1) with a fixed *R*, we applied the alternating least-squares (ALS) method (via the “`cp-als.m`” function from the Tensor Toolbox (www.tensortoolbox.org/)). In Section III-D, we further investigated the impact of *R* on the performance of TKL.

#### 3) Nonlinear graph fusion (NGF)

In this stage, data from different sources are represented as graphs: the nodes are subjects and edges depicts between-subject modalities. Nonlinear graph fusion (NGF) [33] combines these graphs such that key information is integrated in a non-linear way to effectively capture the complex, nonlinear interactions between different data types. NGF is an unsupervised process based on the graph diffusion process (DP). The DP is carried out in *T* iterations. In each iteration, the neighborhood information is represented by a sparse matrix **S**. This matrix takes (k-nearest) neighbor information into consideration with values derived from a weighted matrix **W**(*p, q*) = *w*_*pq*_ (the weight of the edge between vertices *p* and *q*) in each graph and normalized weighted values based on the k-nearest neighbor. The DP in each step states: 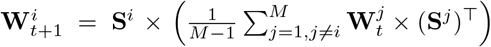 where the superscripts *i* and *j* represent modality indices, the subscript *t* denotes the current iteration, and × indicates matrix multiplication.

#### 4) Kernel Support Vector Machine (SVM)

After TKL and graph diffusion processes, we aggregated kernels from all modalities into a single kernel using summation. To classify subjects, we used a kernel SVM. Kernel functions, such as *k*(**x**_*n*_, **x**_*m*_) = Φ(**x**_*n*_)^⊤^Φ(**x**_*m*_), provide flexibility and enable SVM to handle nonlinear relationships. In linear cases, where 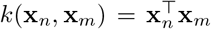, kernel SVM boils down to soft SVM. In this paper, we used kernel matrix **K**(*m, n*) = *k*(**x**_*n*_, **x**_*m*_), which corresponds to the similarity learned from the random forest in the previous step.

Remarkably, we employed semi-supervised learning in our framework by using all data to construct a complete kernel matrix via random forest. The training and testing kernels are then derived from this complete kernel using subject indices. In general, the kernel SVM problem can be expressed as follows:

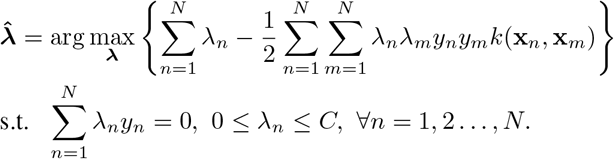

The output/label can be determined as

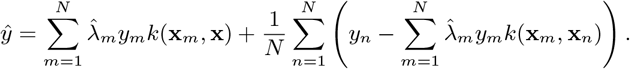

## III. Experiments

We begin by summarising the key findings of the experiments. First, TKL preserves the within-modality structural information and the between-modality complementary information. Second, TKL, combined with NGF, yields clear patterns that produce biologically explainable results. Third, integrating TKL and NGF enhances overall disease classification performance.

Next, we outline the experiments. First, we conducted all experiments using a random holdout method with 80% of the data for training and 20% for testing, repeated 100 times. In other words, each experiment consisted of a random draw and random 80/20 split of the data; there were no overlapping subjects between the training and test sets. To evaluate the kernel matrix for classification, we reported accuracy, specificity, and sensitivity scores and present their means *±* standard errors. For classification purposes, we used the *libsvm* package in MATLAB (www.csie.ntu.edu.tw/~cjlin/libsvm/). For tensor decomposition, we utilized the CP model implemented in the tensor toolbox (www.tensortoolbox.org/), with a tolerance of 10^−9^ and a maximum of 1000 iterations. For NGF, we used five iterations with a neighborhood size of 3.

To compare the kernel/similarity matrix (**K**) of each modality vs the ground truth kernel (**K**_**y**_), we used a scoring method similar to that in [35]. More concretely, the score is defined as follows:

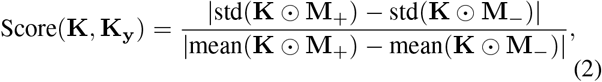

where “ ⊙ “ is the element-wise product, “mean(·)” and “std(·)” denote the mean and standard deviation operations, respectively. With **y** ∈ ℝ^1×*N*^, *y*_*i*_ ∈ *{*−1, 1*}*, then matrix **K**_**y**_ is defined as follows:

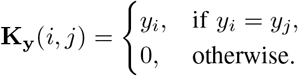

Matrices **M**_+_ and **M**_−_ are binary masks defined as

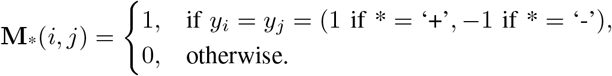

A lower score between the input and the output kernels indicates better separation between individual classes.

### A. TKL preserves underlying structural information

To demonstrate that TKL not only captures the inter-subject relationship but also integrates complementary inter-modality information, we evaluate the similarity scores using Equation (2) between the input and output kernels, both before and after TKL. Table II compares the kernels across three binary pairs of groups – CN vs AD, CN vs MCI, and MCI vs AD – and suggests several interesting findings.

**TABLE II.** Comparision between original kernel matrix and the kernel after TKL with output kernel.

| Group | Modality | Score with output kernel |  |
| --- | --- | --- | --- |
|  |  | Original | TKL |
| CN vs AD | GM | 18.63 ± 0.13 | 9.22 ± 0.12 (↓ 50.56%) |
| | PET | 4.44 ± 0.02 | 3.98 ± 0.06 (↓ 10.36%) |
| | CSF | 16.85 ± 0.25 | 6.24 ± 0.1 (↓ 62.97%) |
| | SNP | 77.95 ± 0.48 | 56.58 ± 0.96 (↓ 27.45%) |
| CN vs MCI | GM | 48.35 ± 0.38 | 46.51 ± 0.47 (↓ 3.80%) |
| | PET | 19.94 ± 0.15 | 17.16 ± 0.13 (↓ 14.02%) |
| | CSF | 72.23 ± 0.49 | 41.00 ± 0.85 (↓ 43.30%) |
| | SNP | 171.16 ± 1.88 | 141.39 ± 1.44 (↓ 17.38%) |
| MCI vs AD | GM | 75.99 ± 0.50 | 68.38 ± 0.70 (↓ 10.02%) |
| | PET | 24.09 ± 0.20 | 20.70 ± 0.20 (↓ 14.08%) |
| | CSF | 69.09 ± 0.45 | 38.96 ± 0.56 (↓ 43.53%) |
| | SNP | 215.22 ± 2.41 | 178.64 ± 2.27 (↓ 17.00%) |

First, overall the classification results show the best for CN vs AD, followed by CN vs MCI, and then MCI vs AD. Intuitively, this makes sense as the difference between CN and AD is larger than that of the other two groups, and MCI is perhaps more similar to AD than to CN. Our results, however, show that TKL detected this from an arithmetic and automated way. Second, in all three groups, PET exhibits the highest performance with mean scores of 4.44, 19.94, and 24.09 for CN vs AD, CN vs MCI, and MCI vs AD, respectively. Third, the results using SNPs data exhibit the lowest performance. Nevertheless, using TKL, all modalities show an improvement with the output kernel. This improvement can be explained by TKL’s ability to use complimentary information between modalities and the underlying rank structure constraint.

Fourth, CSF demonstrates a significant improvement in accuracy across the three calssification tasks (decreasing by 62.97% in CN vs AD, 43.30% in CN vs MCI, and 43.53% in MCI vs AD). Our analyses thus hint that GM, PET, and SNP data may effectively complement CSF in broad scenarios.

Fifth, in CN vs AD groups, the TKL method enhances the score of the GM modality by integrating complementary PET, CSF, and SNP modalities. This integration significantly reduces the error in the output kernel by 50.56% compared to the original kernel. However, in the MCI vs AD groups, it decreases by 10.02%. This indicates that the boundary between multimodal data obtained from MCI vs AD subjects and that obtained from CN vs MCI subjects (and thus the projections from multimodal data to subject groups) are not as clear as those obtained from CN and AD groups. Due to such nebulous (feature)-modality boundaries and less accurate feature-to-label projections, it is more difficult to obtain complementary information from each modality to build as discernible a decision boundary between MCI vs AD groups and CN vs MCI groups as it between CN vs AD groups.

### B. TKL enhances overall classification performance

To provide an overview of TKL’s performance, we conducted experiments across three classification tasks (CN vs AD, CN vs MCI, and MCI vs AD), modality types (single modalities vs combinations (summations)), and methods (original tensor method vs NGF vs TKL + NGF). Table III presents a performance comparison between single modality and multimodal fusion using different fusion strategies. To evaluate the original kernel matrix, we first pre-computed each kernel from results from the random forests. We then fed each modality-specific kernel into a kernel SVM to perform classification. For NGF, we added a DP after obtaining the modality-specific kernel matrix to exchange information across modalities before feeding them into the kernel SVM. Finally, we compared this approach with TKL (see Fig. 3). In TKL + NGF, the original kernel went through TKL to enhance the underlying structure and utilised complementary information before proceeding to the diffusion process to capture neighborhood information. Finally, the terminal kernel matrix was fed into the kernel SVM for classification. There are a few points to note. First, among single modalities, PET exhibits the highest performance in all three classification tasks: CN vs AD, CN vs MCI, and MCI vs AD. This is followed by GM, CSF, and SNP. SNP demonstrates significantly lower performance compared to the other three modalities. These results are in line with the score comparisons discussed in Section III-C. Second, although NGF performs well in some cases for fusing four modalities or in cases with only three modalities (GM, CSF, PET), when considering all four modalities and averaging over 100 experiments, it shows instability and higher error rates compared to using the original kernel and the TKL. When fusing all modalities, in CN vs AD, NGF yields an error rate of 0.65 compared to 0.41 using the original kernel and 0.49 using the TKL kernel. In CN vs MCI, NGF results in an error rate of 0.81 compared to 0.59 and 0.62 for the original and TKL kernels, respectively. For AD vs MCI, NGF results in an error rate of 0.84 compared to 0.48 and 0.21 for the original and TKL kernels, respectively. Third, TKL outperforms the original kernel and NGF, achieving a mean accuracy of 91.31%, 81.45%, and 78.27% for classifying CN vs AD, CN vs MCI, and AD vs MCI, respectively.

**TABLE III.** Performance of single modality vs multimodal fusion under different fusion strategies.

| Group | Modality | Original |  |  | NGF |  |  | NGF+TKL |  |  |
| --- | --- | --- | --- | --- | --- | --- | --- | --- | --- | --- |
| | | Accuracy (↑) | Specificity (↑) | Sensitivity (↑) | Accuracy (↑) | Specificity (↑) | Sensitivity (↑) | Accuracy (↑) | Specificity (↑) | Sensitivity (↑) |
| CN vs AD | GM | 78.81 ± 0.54 | 83.24 ± 0.59 | 89.45 ± 0.99 | 83.24 ± 0.59 | 86.09 ± 0.73 | 80.46 ± 0.95 | 89.45 ± 0.99 | 91.98 ± 0.69 | 87.54 ± 1.94 |
| | PET | 85.48 ± 0.50 | 87.85 ± 0.47 | 89.52 ± 0.91 | 87.85 ± 0.47 | 89.17 ± 0.69 | 86.69 ± 0.84 | 89.52 ± 0.91 | 92.51 ± 0.65 | 86.77 ± 1.90 |
| | CSF | 76.46 ± 0.50 | 75.46 ± 0.57 | 87.92 ± 1.11 | 75.46 ± 0.57 | 78.26 ± 0.92 | 73.06 ± 1.06 | 87.92 ± 1.11 | 92.84 ± 0.66 | 83.49 ± 2.12 |
| | SNP | 49.49 ± 0.71 | 55.47 ± 0.79 | 89.96 ± 0.79 | 55.47 ± 0.79 | 60.27 ± 1.32 | 50.62 ± 1.35 | 89.96 ± 0.79 | 87.95 ± 1.30 | 92.45 ± 0.58 |
| | Combine | 88.72 ± 0.41 | 88.63 ± 0.61 | 88.98 ± 0.74 | 80.10 ± 0.65 | 83.47 ± 1.08 | 76.54 ± 1.32 | 91.31 ± 0.49 | 89.88 ± 0.87 | 93.06 ± 0.50 |
| CN vs MCI | GM | 66.00 ± 0.62 | 73.72 ± 0.74 | 80.39 ± 0.91 | 73.72 ± 0.74 | 78.84 ± 1.12 | 68.53 ± 1.08 | 80.39 ± 0.91 | 83.18 ± 1.31 | 78.14 ± 1.11 |
| | PET | 70.89 ± 0.60 | 78.94 ± 0.59 | 82.50 ± 0.67 | 78.94 ± 0.59 | 82.84 ± 0.73 | 75.20 ± 1.13 | 82.50 ± 0.67 | 86.81 ± 0.80 | 77.81 ± 1.20 |
| | CSF | 60.44 ± 0.57 | 57.89 ± 0.62 | 78.83 ± 0.79 | 57.89 ± 0.62 | 68.11 ± 1.15 | 46.91 ± 1.25 | 78.83 ± 0.79 | 80.51 ± 0.96 | 76.95 ± 1.23 |
| | SNP | 49.50 ± 0.85 | 58.56 ± 0.69 | 80.00 ± 0.74 | 58.56 ± 0.69 | 67.80 ± 1.76 | 49.37 ± 1.94 | 80.00 ± 0.74 | 78.29 ± 1.06 | 82.81 ± 0.99 |
| | Combine | 72.11 ± 0.59 | 79.78 ± 0.79 | 64.26 ± 1.45 | 67.11 ± 0.81 | 70.91 ± 1.92 | 64.86 ± 2.03 | 81.45 ± 0.62 | 80.95 ± 0.91 | 82.80 ± 0.88 |
| AD vs MCI | GM | 60.56 ± 0.97 | 62.05 ± 1.03 | 75.45 ± 0.93 | 62.05 ± 1.03 | 58.65 ± 1.71 | 65.20 ± 1.40 | 75.45 ± 0.93 | 69.77 ± 2.06 | 81.63 ± 0.85 |
| | PET | 73.51 ± 0.51 | 65.62 ± 0.64 | 72.62 ± 1.03 | 65.62 ± 0.64 | 69.51 ± 1.10 | 62.78 ± 1.18 | 72.62 ± 1.03 | 68.34 ± 2.01 | 76.16 ± 1.93 |
| | CSF | 60.71 ± 0.51 | 57.14 ± 0.65 | 73.36 ± 0.93 | 57.14 ± 0.65 | 51.15 ± 1.53 | 63.84 ± 1.26 | 73.36 ± 0.93 | 73.29 ± 1.20 | 73.84 ± 1.72 |
| | SNP | 49.85 ± 0.59 | 52.38 ± 0.68 | 76.93 ± 0.41 | 52.38 ± 0.68 | 38.85 ± 1.73 | 66.11 ± 1.66 | 76.93 ± 0.41 | 74.19 ± 1.48 | 79.24 ± 0.60 |
| | Combine | 71.58 ± 0.48 | 67.10 ± 0.93 | 75.97 ± 0.57 | 57.14 ± 0.84 | 45.16 ± 2.26 | 69.68 ± 1.70 | 78.27 ± 0.21 | 76.86 ± 0.94 | 79.91 ± 0.59 |

### C. TKL provides well-explained results

We have so far demonstrated the utility of the TKL method in classification tasks via kernel learning. Although TKL improves classification performance, since the kenels were derived from biological data, naturally, one would ask “are the kernels biologically explainable”? To investigate TKL’s explainability, we visualized the kernel matrix of the original kernel and the diffusion process (DP) before and after applying TKL in Figure 5. For demonstration purposes, we showed two modalities, the PET and CSF modalities. Our results suggest that, compared to directly applying the DP to the original kernel, TKL derived a clearer and more structured pattern (see Fig. 5). Although the kernel matrix after DP improves classification, it tends to lose structural information and class patterns. The DP in NGF leverages neighborhood information and exchange it among modalities; the significant variance between subjects and modalities, however, may trigger problems when directly interchanging information without normalization. To address this issue, we suggest incorporating TKL before the DP: this not only enhances classification performance but also yields sparse kernel matrices. These matrices can be further analyzed to provide a deeper understanding of the connections between subjects and offer better insights into personalized diagnosis.

**Fig. 5.**
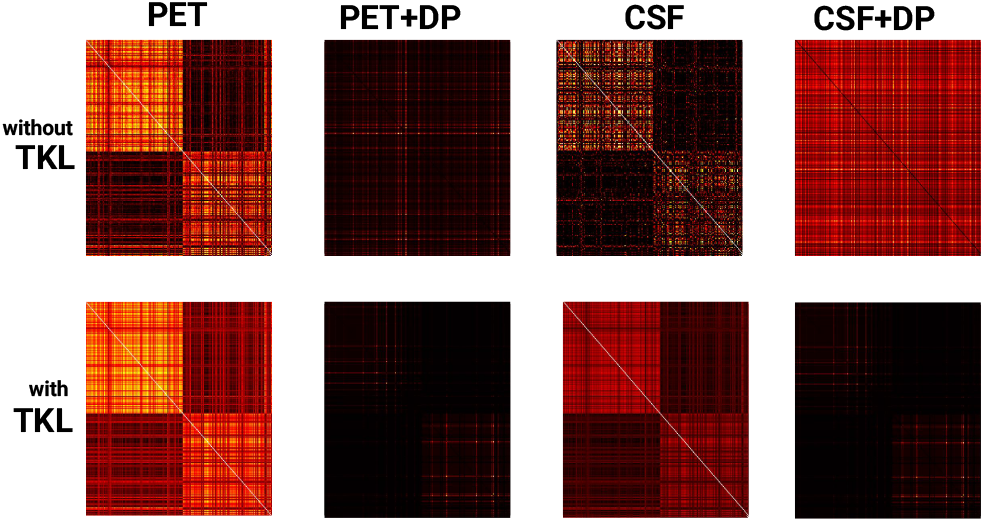
Kernels learned with and without TKL for PET and CSF data.

### D. Influence of rank in TKL

TKL leverages the underlying rank structure, where the rank equals to the number of classes. In binary classification, we set rank to two for all tasks: CN vs AD, CN vs MCI and MCI vs AD. To evaluate how the rank affects the overall error across modalities, we calculated the mean score among the input and output kernels using ranks ranging from 2 to 10 (see Fig. 6).

**Fig. 6.**
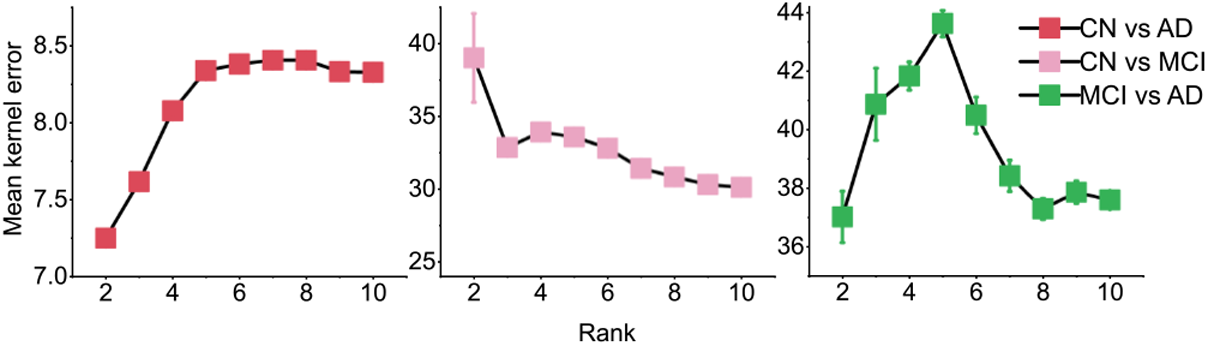
Influence of rank in tensor decomposition.

There are several interesting patterns. First, in CN vs AD classification, the tensor kernel with *R* = 2 yields the smallest error. The error increases until *R* = 6 with a minor drop from *R* = 7 to *R* = 10. Second, in CN vs MCI classification, the tensor kernel with *R* = 2 yields the highest error and high variance. The error decreases as the rank increases. Third, in MCI vs AD classification, the error increases from *R* = 2 to *R* = 5 (similar to the CN vs AD pattern) and decreases from *R* = 5 to *R* = 10 (similar to the CN vs MCI pattern).

## IV. Conclusion

We introduced TKL to fuse multi-modal data for classification. By constructing a low-rank tensor, TKL learns the relevant but succinct underlying structure and complementary information from multi-modal data. It is robust to high variance among modalities, enhances pattern recognition, and improves classification performance. Experiments using the ADNI data across three classification tasks show that TKL outperforms both kernel approaches with linear combinations and nonlinear combinations with diffusion processes. Despite TKL’s advantage in classification and kernel pattern recognition, it omits feature selections, as the method operates at the group-level kernel matrices. Identifying key features before data fusion may be useful in biomarker studies in medical research. Future studies could explore the use of tensor decomposition for both feature selection and data fusion.

## References

[1] C. L. Masters, R. Bateman, K. Blennow, C. C. Rowe, R. A. Sperling, and J. L. Cummings, “Alzheimer’s disease,” Nat. Rev. Dis. Primers, vol. 1, no. 1, pp. 1–18, 2015.

[2] Alzheimer’s Disease International, “World Alzheimer Report 2023: Reducing Dementia Risk: Never too early, never too late,” 2023.

[3] Z. Breijyeh and R. Karaman, “Comprehensive Review on Alzheimer’s Disease: Causes and Treatment,” Molecules, vol. 25, no. 24, p. 5789, Dec. 2020.

[4] “2023 Alzheimer’s disease facts and figures,” Alzheimers Dement., vol. 19, no. 4, pp. 1598–1695, 2023.

[5] J. M. Long and D. M. Holtzman, “Alzheimer Disease: An Update on Pathobiology and Treatment Strategies,” Cell, vol. 179, no. 2, pp. 312–339, Oct. 2019.

[6] C. R. Jack and D. M. Holtzman, “Biomarker Modeling of Alzheimer’s Disease,” Neuron, vol. 80, no. 6, pp. 1347–1358, Dec. 2013.

[7] M. Tanveer, B. Richhariya, R. U. Khan, A. H. Rashid, P. Khanna, M. Prasad, and C. T. Lin, “Machine Learning Techniques for the Diagnosis of Alzheimer’s Disease: A Review,” ACM Trans. Multimedia Comput. Commun. Appl., vol. 16, no. 1s, pp. 1–35, Jan. 2020.

[8] V. Berti, R. Osorio, L. Mosconi, Y. Li, S. De Santi, and M. De Leon, “Early Detection of Alzheimer’s Disease with PET Imaging,” Neurodegenerative Diseases, vol. 7, no. 1-3, pp. 131–135, 2010.

[9] L. Mosconi, R. Mistur et al., “FDG-PET changes in brain glucose metabolism from normal cognition to pathologically verified Alzheimer’s disease,” European Journal of Nuclear Medicine and Molecular Imaging, vol. 36, no. 5, pp. 811–822, May 2009.

[10] G. e. a. Chételat, “Amyloid-PET and 18F-FDG-PET in the diagnostic investigation of Alzheimer’s disease and other dementias,” Lancet Neurol., vol. 19, no. 11, pp. 951–962, Nov. 2020.

[11] K. Blennow and H. Hampel, “CSF markers for incipient Alzheimer’s disease,” Lancet Neurol, vol. 2, no. 10, pp. 605–613, Oct. 2003.

[12] Y.-D. Zhang, Z. Dong et al., “Advances in multimodal data fusion in neuroimaging: Overview, challenges, and novel orientation,” Inf.Fusion, vol. 64, pp. 149–187, 2020.

[13] N. Luo, W. Shi, Z. Yang, M. Song, and T. Jiang, “Multimodal fusion of brain imaging data: Methods and applications,” Mach. Intell. Res., vol. 21, no. 1, pp. 136–152, 2024.

[14] J. Sui et al., “Three-way (n-way) fusion of brain imaging data based on mCCA+ jICA and its application to discriminating schizophrenia,” NeuroImage, vol. 66, pp. 119–132, 2013.

[15] S. Qi et al., “Three-way parallel group independent component analysis: Fusion of spatial and spatiotemporal magnetic resonance imaging data,” Human Brain Mapping, vol. 43, no. 4, pp. 1280–1294, 2022.

[16] S. Dhifallah, Rekik, ADNI et al., “Clustering-based multi-view network fusion for estimating brain network atlases of healthy and disordered populations,” J. Neurosci. Methods, vol. 311, pp. 426–435, 2019.

[17] A. Riaz, M. Asad, E. Alonso, and G. Slabaugh, “Fusion of fMRI and non-imaging data for ADHD classification,” Comput. Med. Imaging Graph., vol. 65, pp. 115–128, 2018.

[18] D. Zhang and D. Shen, “Multi-modal multi-task learning for joint prediction of multiple regression and classification variables in Alzheimer’s disease,” NeuroImage, vol. 59, no. 2, pp. 895–907, 2012.

[19] L. Xiao, J. M. Stephen, T. W. Wilson, V. D. Calhoun, and Y.-P. Wang, “A manifold regularized multi-task learning model for iq prediction from two fMRI paradigms,” IEEE Trans. Biomed. Eng., vol. 67, no. 3, pp. 796–806, 2019.

[20] M. Gönen and E. Alpaydın, “Multiple kernel learning algorithms,” J. Mach. Learn. Res., vol. 12, pp. 2211–2268, 2011.

[21] L. He, C.-T. Lu, H. Ding, S. Wang, L. Shen, P. S. Yu, and A. B. Ragin, “Multi-way multi-level kernel modeling for neuroimaging classification,” in Proc. IEEE Conf. Comput. Vis. Pattern Recogn., 2017, pp. 356–364.

[22] Y. Panagakis, J. Kossaifi, G. G. Chrysos, J. Oldfield, M. A. Nicolaou, A. Anandkumar, and S. Zafeiriou, “Tensor methods in computer vision and deep learning,” Proc. IEEE, vol. 109, no. 5, pp. 863–890, 2021.

[23] A. Shashua and T. Hazan, “Non-negative tensor factorization with applications to statistics and computer vision,” in Proc. Int. Conf. Mach. Learn., 2005, pp. 792–799.

[24] J. C. Ho, J. Ghosh, and J. Sun, “Marble: high-throughput phenotyping from electronic health records via sparse nonnegative tensor factorization,” in Proc. ACM SIGKDD Int. Conf. Knowl. Disc. Data Mining, Aug. 2014, pp. 115–124.

[25] S. Ramdhani, E. Navarro, E. Udine, A. G. Efthymiou, B. M. Schilder, M. Parks, A. Goate, and T. Raj, “Tensor decomposition of stimulated monocyte and macrophage gene expression profiles identifies neurodegenerative disease-specific trans-eQTLs,” PLOS Genet., vol. 16, no. 2, p. e1008549, Feb. 2020.

[26] H. Becker, L. Albera, P. Comon, M. Haardt, G. Birot, F. Wendling, M. Gavaret, C. G. Bénar, and I. Merlet, “EEG extended source localization: Tensor-based vs. conventional methods,” NeuroImage, vol. 96, pp. 143–157, Aug. 2014.

[27] T. T. Le, K. Abed-Meraim, P. Ravier, O. Buttelli, and A. Holobar, “Tensor decomposition meets blind source separation,” Signal Process., vol. 221, p. 109483, 2024.

[28] N. D. Sidiropoulos, L. De Lathauwer, X. Fu, K. Huang, E. E. Papalexakis, and C. Faloutsos, “Tensor decomposition for signal processing and machine learning,” IEEE Trans. Signal Process., vol. 65, no. 13, pp. 3551–3582, 2017.

[29] L. T. Thanh, K. Abed-Meraim, N. L. Trung, and A. Hafiane, “A contemporary and comprehensive survey on streaming tensor decomposition,” IEEE Trans. Knowl. Data Eng., vol. 35, no. 11, pp. 10 897–10 921, 2022.

[30] D. Lahat, T. Adali, and C. Jutten, “Multimodal Data Fusion: An Overview of Methods, Challenges, and Prospects,” Proc. IEEE, vol. 103, no. 9, pp. 1449–1477, Sep. 2015.

[31] L. T. Thanh, N. T. A. Dao, N. V. Dung, N. L. Trung, and K. Abed-Meraim, “Multi-channel EEG epileptic spike detection by a new method of tensor decomposition,” J. Neural Eng., vol. 17, no. 1, p. 016023, 2020.

[32] K. R. Gray, P. Aljabar, R. A. Heckemann, A. Hammers, and D. Rueckert, “Random forest-based similarity measures for multi-modal classification of Alzheimer’s disease,” NeuroImage, vol. 65, pp. 167–175, Jan. 2013.

[33] T. Tong, K. Gray, Q. Gao, L. Chen, and D. Rueckert, “Multi-modal classification of Alzheimer’s disease using nonlinear graph fusion,” Pattern Recogn., vol. 63, pp. 171–181, Mar. 2017.

[34] A. Schaefer, R. Kong, E. M. Gordon, T. O. Laumann, X.-N. Zuo, A. J. Holmes, S. B. Eickhoff, and B. T. Yeo, “Local-global parcellation of the human cerebral cortex from intrinsic functional connectivity MRI,” Cerebral Cortex, vol. 28, no. 9, pp. 3095–3114, 2018.

[35] C. H. Nguyen and T. B. Ho, “An efficient kernel matrix evaluation measure,” Pattern Recogn., vol. 41, no. 11, pp. 3366–3372, Nov. 2008.

